# Rdh54 stabilizes Rad51 at displacement loop intermediates to regulate genetic exchange between chromosomes

**DOI:** 10.1101/2022.05.19.492663

**Authors:** Margaret Keymakh, Jennifer Dau, Bryan Ferlez, Michael Lisby, J. Brooks Crickard

## Abstract

Homologous recombination (HR) is a double-strand break DNA repair pathway that preserves chromosome structure. To repair the damaged recipient DNA, HR requires an intact donor DNA sequence located elsewhere in the genome. After the double-strand break is repaired, DNA sequence information can be transferred between donor and recipient DNA molecules through different mechanisms, including DNA crossovers that form between homologous chromosomes. Regulating this transfer of information is an important step in effectively completing HR and maintaining genome integrity. For example, mitotic exchange of information between homologous chromosomes can result in loss-of-heterozygosity (LOH) in diploid organisms, and in higher eukaryotes, the development of cancer. The DNA motor protein Rdh54 is a highly conserved DNA translocase that functions during HR but has a limited role in repairing DNA. Instead, several existing phenotypes in *rdh54Δ* strains suggest that Rdh54 may regulate the flow of information between donor and recipient DNA molecules post DNA repair. In our current study, we used a combination of biochemical and genetic techniques to dissect the role of Rdh54 on the exchange of genetic information after DNA repair. Our data indicates that *RDH54* regulates DNA sequence exchange between chromosomes by limiting the disruption of Rad51 at an early HR intermediate called the displacement loop (D-loop). Rdh54 also protects Rad51 filaments, acting in opposition to Rad51 removal by the DNA motor protein Rad54. Furthermore, we find that expression of a catalytically inactivate allele of Rdh54, *rdh54K318R*, displays a different distribution of information exchange outcomes than *rdh54*Δ cells. From these results, we propose a model for how Rdh54 may effectively regulate information transfer during homologous recombination.

## Introduction

Homologous recombination (HR) is a universally conserved DNA double-strand break repair (DSBR) pathway that functions to maintain chromosome integrity. Failures and mis-regulation of HR can lead to the loss of genome integrity and the development of human cancers [1, 2]. In eukaryotes, HR proceeds through a series of defined steps that begin with the identification of double-strand breaks and the resection of dsDNA into tracts of ssDNA that are bound by replication protein A (RPA). During mitotic DNA repair in eukaryotes, RPA is replaced by the recombinase Rad51 [3]. An alternate name for the ssDNA bound by Rad51 is the recipient DNA, as it will receive information from another DNA molecule during the HR reaction. Rad51 then facilitates the recruitment of the DNA motor protein Rad54 to promote a systematic search of the genome for a homologous region of DNA [4]. Identification of a homologous stretch of DNA results in the formation of a displacement loop (D-loop) intermediate [5-8]. The identified sequence of homologous DNA can also be referred to as the donor DNA, because it donates information to the HR reaction.

D-loops are major intermediates in the HR pathway [9, 10]. They are reversible three strand DNA structures that form between the donor and recipient DNA molecules. D-loop stability is regulated by efficiency of DNA base pairing [11], and the activity of specific helicases [5, 9, 12], topoisomerases [13], and translocases [14, 15]. In budding yeast D-loop reversal is believed to be controlled by the highly conserved DNA motor proteins Mph1, Sgs1-Top3-Rmi1 (STR), Srs2, and Rad54. In each case, protein occupancy at D-loops is an important regulator of reversibility. For example, Mph1 and STR can disrupt D-loops occupied by Rad54 [13, 16], whereas Srs2 cannot [16]. These observations highlight that the timing of protein occupancy at HR intermediates often regulates the rate of downstream recombination outcomes. D-loop reversibility provides opportunities for quality control to ensure that a correct site of homology has been identified and is often balanced with the process of donor DNA target commitment [5]. Current models implicate Rad54 in the removal of Rad51 from the 3’ end of newly stabilized D-loops, thus creating a momentary instability that can be stabilized by the activity of DNA synthesis factors PCNA, RFC, and DNA Polymerase delta [15, 17-19]. Functionally, these factors work to extend the D-loop and restore the information that was lost with the original dsDNA break.

After DNA extension, D-loops can mature into four-strand DNA intermediates through Rad52 mediated annealing of the second end of DNA into an active extended D-loops [20-23], promoting the classical double-strand break repair pathway (DSBR) [8]. Alternatively, extended D-loops can be disrupted and then annealed to the second end of the broken DNA molecule allowing for fill in and completion of DNA repair as part of the synthesis dependent strand annealing (SDSA) [1, 24-26]. If the second end of DNA is not located, the helicase Pif1 can join the DNA replication machinery to create a migrating bubble that can synthesize long tracts of DNA in a process called break induced replication (BIR) [27-29]. The net result of these pathways is non-crossover gene conversion (NCO), gene conversion with crossovers (CO), and gene conversion with BIR. The later of these two outcomes can result in the transfer of information between the recipient and donor DNA molecules. When different chromosomes are used for DNA repair this can lead to allelic loss within the genome, otherwise known as loss-of-heterozygosity (LOH), and is a common cause of certain types of cancers [30].

Rdh54 (Rad54 homolog, a.k.a. Tid1) [31] is a paralog of Rad54 and they share 41% sequence identity [12]. Both proteins are ATP dependent dsDNA translocases and have similar biochemical activities *in vitro*, including DNA supercoiling [32-34], nucleosome remodeling [35, 36], enhancement of Rad51 mediated D-loop formation [4, 32, 33, 37, 38], and movement along dsDNA [39, 40]. Several biochemical differences have also been observed between Rdh54 and Rad54. These include the observation that Rad54 preferentially initiates Rad51 mediated homology search [4, 41], that Rdh54 stabilizes Rad51 filaments on ssDNA against ATP depletion, and that each translocase shares a unique binding site on Rad51 filaments [42]. Major differences in *RDH54* and *RAD54* activity manifests in cells during HR. For example, *RAD54* is required for HR mediated gene conversion during repair of damaged DNA in *Saccharomyces cerevisiae*, whereas *RDH54* is not [12, 43]. In contrast, *RDH54* is required for inter-chromosomal template switching (ICTS), but not simple gene conversion [7]. ICTS events are when D-loops move between chromosomes and are associated with gene conversion with BIR outcomes [44-46]. Despite these well characterized differences, both motor proteins are implicated in the process of D-loop expansion [9, 15], where their mechanisms remain poorly defined. Moreover, the role of Rdh54 during post gene conversion DNA repair steps also remains an open question, with possibilities including the regulation of CO to NCO outcomes, BIR transitions, or DNA second end capture.

Here we have conducted a study on the biochemical and genetic mechanism by which Rdh54 effects downstream HR outcomes through regulation of D-loop intermediates. We conclude that Rdh54 can act as a physical barrier to Rad54 at D-loop intermediates in a position dependent manner. The role of this interaction is to regulate BIR. We also find that loss of Rdh54 catalytic activity results in a dramatic reduction in mitotic crossover outcomes. These data suggest that Rdh54 must be removed to grant access to pathways that result in mitotic crossover outcomes. Finally, we present a model for how Rdh54 may function as a mobile barrier during HR to regulate the exchange of information between recipient and donor DNA molecules.

## Results

### Deletion of Rdh54 results in an increase in BIR HR outcomes

To initiate our studies on the role of *RDH54* in HR outcomes, we used a defined genetic assay to monitor HR outcomes between homologous chromosomes [47]. We used this assay for two reasons. First, *RDH54* has been shown to have greater influence on HR between homologous chromosomes [43]. Second, this assay would allow us to understand how the population of HR outcomes were influenced by *RDH54*. Therefore, even if only a fraction of cells utilized *RDH54* post DNA repair, we would still be able to observe changes to this specific population. This assay relies on an I-SceI nuclease site located in the ORF of an inactive *ADE2* (*ade2-I*) gene located on chromosome XV (Fig. 1A). Downstream of the I-SceI cut site is an *HPHMX* marker. The homologous chromosome contains a non-cleavable, but also inactive, allele of *ADE2* (*ade2-n*), and a *NATMX* marker located downstream of *ade2-n* (Fig 1A). The differences in the *ADE2* alleles allow for one chromosome to act as a substrate for the I-SceI nuclease, and the other homolog serves as an uncut template for repair. Importantly, *URA3* and *MET22* markers located on the far end of chromosome XV act as markers to report on loss of chromosomes (Fig 1A).

**Figure 1:**
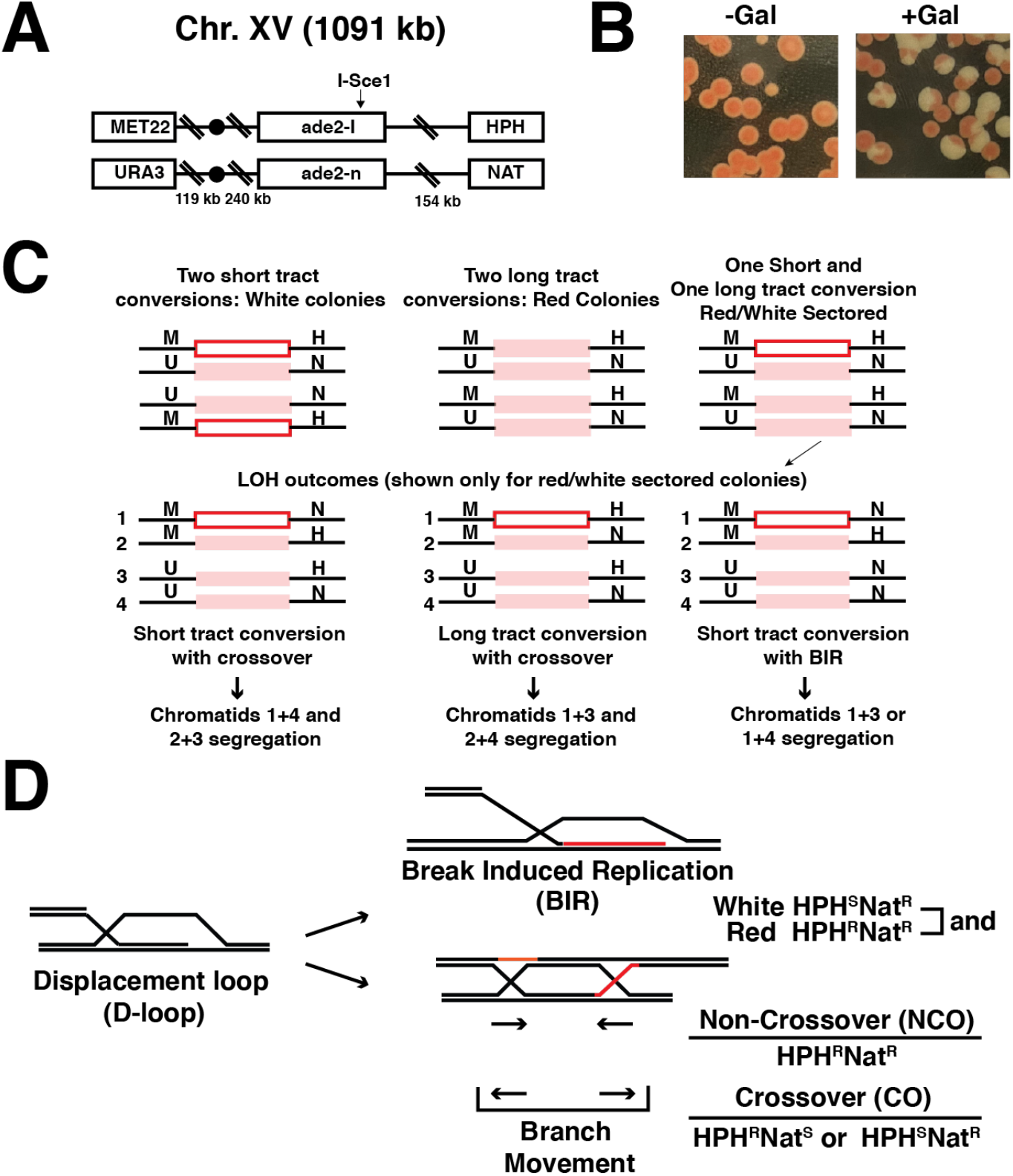
Analysis of RDH54 in post gene conversion recombination. **(A)**. Schematic diagram illustrating assay used to monitor recombination between homologous chromosomes during a single double strand DNA break. The reporter is located on chromosome XV and only one homolog has an active I-SceI site **(B)**. Yeast colonies plated after ± induction of I-SceI using galactose. **(C)**. Schematic diagram illustrating the potential outcomes for DNA repair via homologous recombination. In the first step gene conversion can occur through long tract or short tract DNA repair. Following cellular division, non-crossover, crossover, and break induced replication (non-reciprocal exchange) outcomes can be determined for each event. **(D)**. Cartoon illustration of HR outcomes that can be determined from this assay.

When galactose is added to the media, overexpression of I-SceI results in the cleavage of ∼70% of both sister chromatids [47, 48] and forces DNA repair using the homologous chromosomes. DNA repair by gene conversion results in the formation of white colonies (Short tract GC), red colonies (Long tract GC), and sectored colonies (Fig 1BC). Sectored colonies are the result of cells that undergo division after plating and are the most useful in determining HR outcomes (Fig 1C). Therefore, in our experiments we only measured HR outcomes for sectored colonies. The different antibiotic markers located on either homolog allows for the interpretation of HR outcomes. For example, cells that have undergone long or short tract gene conversion with a crossover will have chromatid segregants that result in LOH (Fig 1CD). LOH results in the formation of colonies in which one sector is resistant to either hygromycin (HPH) or nourseothricin (clonNAT), but not both. LOH can also result from BIR. Sectored colonies that have gone through BIR will have a white sector that is resistant to clonNAT, but not HPH. This results from the preferential replication of one sister chromatid due to failed second end capture [29, 49, 50]. Sectored colonies in which both sectors are resistant to HPH and clonNAT are the results of non-crossover DNA repair outcomes (Fig 1CD).

We first measured the number of cells that survived DNA repair after I-SceI cleavage by counting the number of colonies that grew from cultures treated with or without galactose. By dividing the number of colonies from ± galactose plates we were able to calculate a plating efficiency. The plating efficiency of WT cells was 77±3.3% (Fig 2A). Similar values were observed for *RDH54/rdh54Δ* (83±4.4%), *rdh54Δ/rdh54Δ* (69±4.1%), *RDH54-KanMX/RDH54-KanMX* (83.3± 10.6%), and *rdh54K318R/rdh54K318R* (an ATPase inactive allele of Rdh54) (75.5±8%) strains (Fig 2A). In contrast, *rad54Δ/rad54Δ* had a plating efficiency of only 34±2.8%, suggesting that cells failed to repair the I-SceI induced DSB (Fig 1D). The plating efficiency for *rad54Δ/rad54Δ* cells was comparable to the percentage of cells that were not cut by I-SceI based on reinduction assays (30±10% uncut in WT). Importantly, *rad54Δ/rad54Δ* failed to produce any sectored or white colonies indicating no recombination had taken place (Supplemental Table S1) and served as an effective negative control. In contrast, WT, *RDH54/rdh54Δ, rdh54Δ/rdh54Δ, RDH54/RDH54-KanMX*, and *rdh54K318R/rdh54K318R* strains all produced white and sectored colonies at similar frequencies upon induction of I-SceI (Fig 2A). Taken together, these data suggest that simple gene conversion and completion of DNA repair occur with similar efficiency in WT and *rdh54* mutant strains.

**Figure 2:**
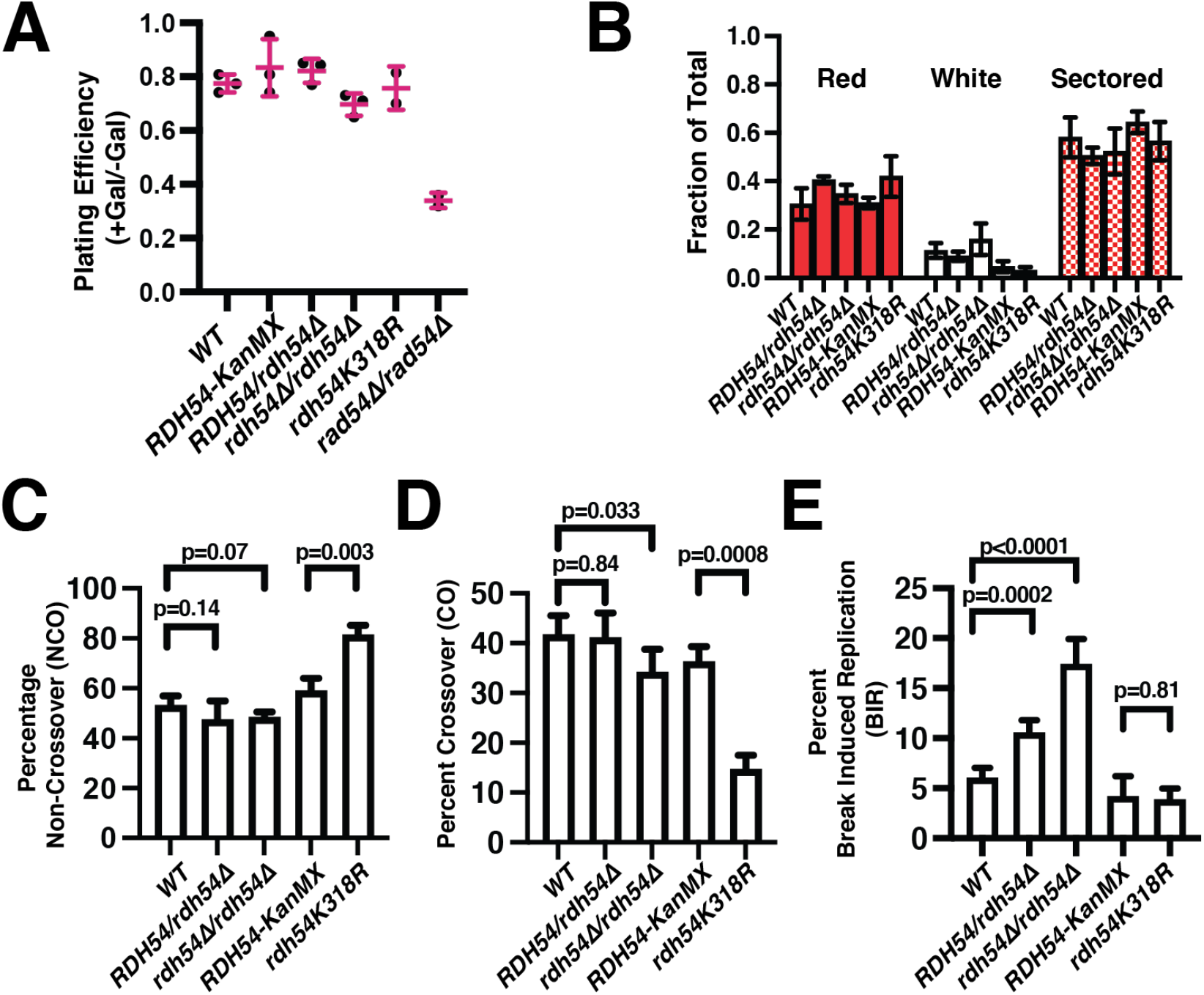
Loss of *RDH54* results in an increase in BIR outcomes. **(A)**. Graph representing the plating efficiency of WT, *RDH54-KanMX/RDH54-KanMX, RDH54/rdh54*Δ, *rdh54Δ/rdh54Δ, rdh54K318R/rdh54K318R*, and *rad54Δ/rad54Δ*. Crossbar and error bars represent the mean and standard deviation of at least 3 independent measurements. **(B)**. Graphical representation of the fraction of cells that are red, white, or sectored after DNA double strand break repair. Strains represented here are *WT, RDH54/rdh54Δ, rdh54Δ/rdh54Δ, RDH54-KanMX/RDH54-KanMX, rdh54K318R/rdh54K318R*. The column and the error bars represent the mean and standard deviation of at least three independent experiments. **(C)**. Graph representing the percentage of non-crossover outcomes for WT, *RDH54-KanMX/RDH54-KanMX, RDH54/rdh54*Δ, *rdh54Δ/rdh54Δ, rdh54K318R/rdh54K318R*. Measurements represent the mean and standard deviation of four independent experiments. **(D)**. Graph representing the percentage of crossover outcomes for WT, *RDH54-KanMX/RDH54-KanMX, RDH54/rdh54*Δ, *rdh54Δ/rdh54Δ, rdh54K318R/rdh54K318R*. Measurements represent the mean and standard deviation of four independent experiments. **(E)**. Graph representing the percentage of non-reciprocal exchange/Break induced replication (BIR) outcomes for WT, *RDH54-KanMX/RDH54-KanMX, RDH54/rdh54*Δ, *rdh54Δ/rdh54Δ, rdh54K318R/rdh54K318R*. Measurements represent the mean and standard deviation of at least four independent experiments.

We next calculated HR outcomes for each of these strains. We found that in WT cells, NCO, CO, and BIR occurred at frequencies of 53.2±3.6%, 41.7±3.7%, and 6.0±1%, respectively (Fig 2CDE). The corresponding frequencies observed for *RDH54/rdh54Δ* cells were 47.4±7.5%, 41.1±5.0%, and 10.5±1.2%, and for *rdh54Δ/rdh54Δ* cells 48.3±2.1%, 34.2±4.6%, 17.4±2.5%, respectively (Fig 2CDE). The critical difference is in the amount of BIR observed between *WT, RDH54/rdh54Δ*, and *rdh54Δ/rdh54Δ* strains, which contain two, one, or no copies of WT Rdh54, respectively. We next complemented the phenotype by integrating WT copies of *RDH54* into the native *RDH54* locus. The integrands had a *KanMX* marker located 125 bp downstream of the end of the *RDH54* ORF. These strains are denoted as *RDH54-KanMX/RDH54-KanMX*. This served as a complementing control as well as a control for other *RDH54* mutant alleles. *RDH54/RDH54-KanMX* had a distribution of 59.4±5.9%, 36.2±3.2%, 4.2±2.1% for the frequency of observed NCO, CO, and BIR outcomes, respectively. While there was some variation in the NCO and CO populations, they were not significantly different from WT. Importantly, this construct did complement the BIR phenotype (Fig 2CDE).

We next asked if ATP hydrolysis was required for inhibition of BIR by measuring outcomes for *rdh54K318R/rdh54K318R*, a catalytically inactive version of *RDH54. rdh54K318R/rdh54K318R* strains resulted in 81.4±3.9%, 14.6±2.8%, and 3.8±1.2%, of NCO, CO, and BIR outcomes, respectively (Fig 2CDE). We were surprised to see that while *rdh54K318R/rdh54K318R* complemented the BIR phenotype, it also resulted in a significant loss (p= 0.0008) of CO outcomes in favor of an increase in NCO outcomes (p=0.0036) (Fig 2BC). This observation was unexpected, but consistent with a previous observation made in haploid yeast strains [9]. A novel finding from these data is that the increase of NCOs and loss of COs do not result in cell death, implying that the *rdh54K318R* mutant yields a trapped but recoverable recombination intermediate in diploid cells (Fig 1D).

To ensure that these outcomes were due to HR and not the loss of chromosomes, sectored colonies were scored for viability on -Ura/-Met plates. Colonies that retain both homologous chromosomes will grow, and if chromosome loss has taken place colonies will fail to grow. No significant increase in chromosome loss was observed for all strains tested (Supplemental Table S1). From these data we conclude that *RDH54* limits BIR in a dosage dependent manner, and failure of the Rdh54 ATPase results in loss of access to gene conversion with crossovers.

### Rdh54 prevents Rad54 mediated D-loop turnover

Current models predict that Rad54 can remove Rad51 from D-loop intermediates, promoting the first steps in gene conversion [15, 17, 24]. Our genetic observations led us to test whether Rdh54 could limit Rad54 mediated D-loop turnover *in vitro*. D-loop reactions can be modelled *in vitro* by performing time course experiments of D-loop formation mediated by Rad51, Rad54 and RPA [15]. D-loop turnover is measured by first observing the formation of D-loops, and then monitoring D-loop disappearance. We used an established *in vitro* system to measure D-loop formation and turnover relying on fluorescently labeled recipient DNA (Atto 647N) composed of a 35 nt non-homologous duplex region with a 21 nt 3’ssDNA extension to serve as a binding site for a single turn of a Rad51 filament (Fig. 3A). Supercoiled pUC19 plasmid served as the donor DNA molecule in these assays. In our experiments the D-loop is considered only the initial strand invasion intermediate and is detected by a gel shift in which the recipient DNA migrates higher in the gel due to an interaction with the donor plasmid. In our hands D-loop disappearance occurs between the 10-minute and 45-minute time points of the reaction (Fig. 3AB).

**Figure 3:**
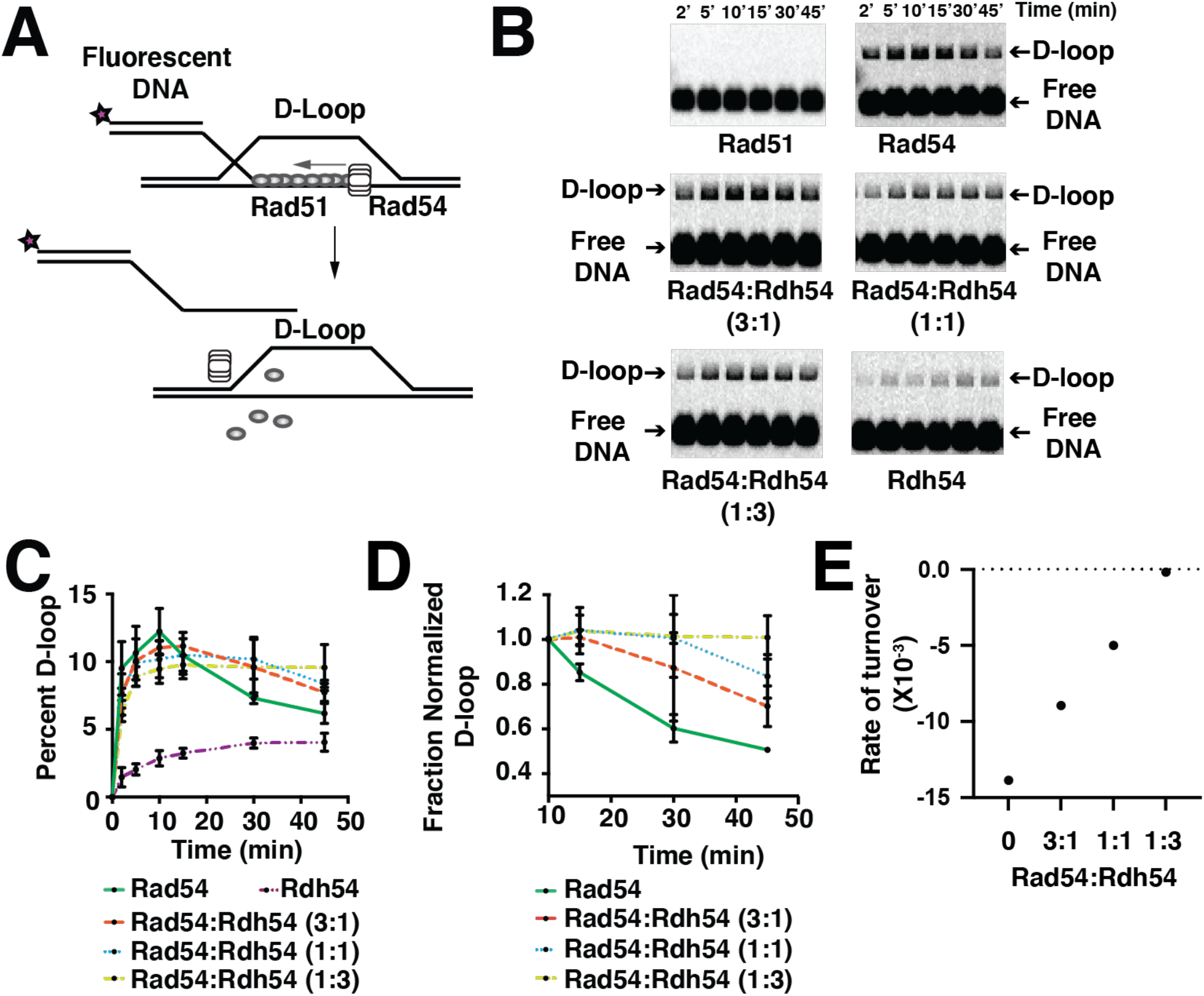
Rdh54 inhibits Rad54 mediated D-loop turnover. **(A)**. Cartoon diagram illustrating the experimental design for monitoring Rad54 mediated D-loop turnover. **(B)**. Representative agarose gels illustrating D-loop turnover for Rad51 alone, Rad51+Rad54 (30 nM), Rad51+Rad54+Rdh54 (10 nM), Rad51+Rad54+Rdh54 (30 nM), Rad51+Rad54+Rdh54 (90 nM), Rad51+Rdh54 (30 nM). **(C)**. Quantification of D-loop turnover experiment as a Fraction of D-loop. The error bars represent the standard deviation of three independent experiments. **(D)**. Normalized quantification of D-loop turnover. Experiments were normalized by setting the maximum fraction D-loop (10 min) to one and normalizing the other time points to that value. The error bars represent the standard deviation of three independent experiments. **(E)**. Graphical depiction of linear fits of the D-loop turnover phase (10-45 min) of the reaction. The rates are represented from the slope of the best fit line for 0, 3:1, 1:1, and 1:3 molar ratios of Rad54:Rdh54.

In this assay the addition of Rad51 alone is not sufficient for D-loop formation. The addition of Rad54 together with Rad51 initially catalyzed the formation of D-loops followed by D-loop turnover (Fig. 3BC). The reaction was monitored at 2, 5, 10, 15, 30, and 45 min, and as expected D-loops formed in the first 10 minutes of the reaction and then began to disappear between 10 minutes and 45 minutes, with 50% of the maximum number of D-loops formed lost after 45 min (Fig 3BC). We next formed D-loop reactions with a fixed concentration of Rad51,Rad54, and RPA and titrated Rdh54 creating molar ratios of 1:3, 1:1, and 3:1 Rad54:Rdh54. Rdh54 in combination with Rad54 had little effect on the formation of D-loops with only a modest reduction of D-loop formation when Rdh54 was present at 3-fold molar excess to Rad54 (Fig 3BC). Surprisingly, a reduction in the D-loop turnover phase of the reaction was observed with the fraction of D-loop disappearance increasing from 0.5 ± 0.1, 0.7± 0.1, 0.83± 0.1, and 1.0±0.1 at 0, 3:1, 1:1, 1:3 molar ratios of Rad54:Rdh54, respectively. This represented a 29%, 39%, and 100% reduction in turnover (Fig 3CDE). Rdh54 in combination with only Rad51 was able to catalyze D-loop formation but did not promote D-loop dissociation within our observation window (Fig 3BCD).

To determine if inhibition was due to the ability of Rdh54 to inhibit the ATPase activity of Rad54, we mixed Rdh54K318R with Rad54 in equal molar amounts in the presence or absence of Rad51. We observed that there was no change in the Rad54 ATPase activity in either case, suggesting the ATPase of Rad54 was unaffected (Supplemental Fig 1A). In an effort to determine if our ratios of Rad54:Rdh54 fell within the physiological range, we measured the total number of Rad54 and Rdh54 molecules per cell [51]. Our measurements focused on the ratio of Rad54 and Rdh54 in diploid cells. We found that Rad54 occurred at a mean 430 molecules per cell and Rdh54 occurred at 673 molecules per cell (Supplemental Fig 1B). This gives a predicted ratio of Rdh54:Rad54 of approximately 1.6:1, suggesting that our *in vitro* experiments were performed around the physiological ratio of the two translocases. Together these data suggest that Rdh54 can prevent D-loop turnover under physiological protein concentrations.

### Rdh54 prevents Rad54 mediated accessibility of the 3’ end at D-loops

The net result of Rad54 activity at D-loops is the accessibility of the 3’ end of DNA to act as a primer for DNA polymerase extension [17]. This can be modelled *in vitro* using Klenow fragment (exo-) as a DNA polymerase to form extended D-loops (Fig 4A). We next asked whether Rdh54 could limit the accessibility of the 3’ end of newly formed D-loop intermediates. The formation of extended D-loops was monitored by first allowing D-loop formation for ten minutes followed by the addition Klenow (exo-) and dNTPs. D-loop extension is detected by a shift in the size of the D-loop within the gel (Fig 4B). No D-loops or shifted products formed in the absence of Rad54 (Fig 4B). When Rad54 was present, extended D-loops represented 17.1±5.8%, 29.5±7.5%, 35.2±5.3% of D-loop populations at 20, 30, and 45 minutes, respectively (Fig 4BC). In contrast, the addition of Rdh54 resulted in the formation of D-loops but did not yield extension products in the time window of the experiment (Fig 4BC). When Rad54 and Rdh54 were mixed, the formation of extended D-loops was inhibited almost completely, with only a small fraction of extended D-loop, 1.4±2.4%, detected at the 45 min time point (Fig 4BC). We do not believe this result is due to the formation of a mixture of Rad54 mediated D-loops and Rdh54 mediated D-loops, as this would result in only a 50% reduction in the number of extended D-loops formed, which is not consistent with our observation. We cannot explicitly rule out the possibility that Rdh54 is blocking Klenow accessibility to the 3’ end. However, together with the result that Rdh54 limits Rad54 mediated D-loop turnover, we are confident that these data suggest Rdh54 is inhibiting Rad54 activity at D-loops.

**Figure 4:**
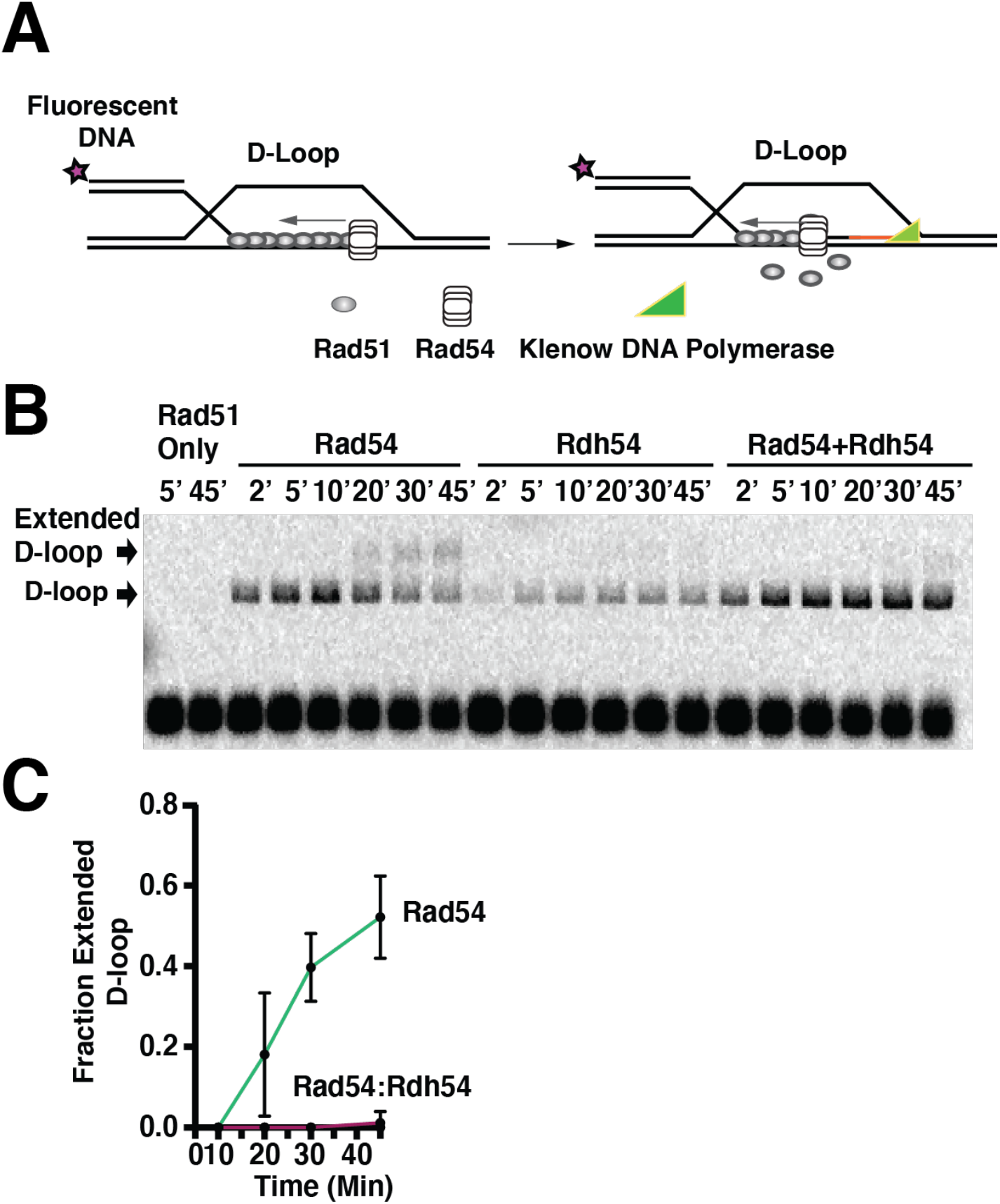
Rdh54 prevents Rad54 from creating accessible 3’ end at D-loop intermediates. **(A)**. Cartoon diagram illustrating the experiment to monitor the 3’ end accessibility by using Klenow (exo-) to extend the nascent D-loop. **(B)**. Representative gel illustrating D-loop extension for Rad51 alone, Rad51+Rad54 (30 nM), Rad51+Rdh54 (30 nM), and Rad51+Rad54 (30 nM) +Rdh54 (30 nM). **(C)**. Quantification of extended D-loop fraction. The error bars represent the standard deviation for three independent experiments.

### The activity of Rdh54 at D-loops is affected by binding position

We next asked whether the unique binding sites of Rad54 and Rdh54 within Rad51 filaments contributed to regulation of D-loop turnover. Previous studies have shown that Rdh54 and Rad54 bind to unique sites within the Rad51 filaments [41]. Unique binding sites on the Rad51 filament were identified by creating chimeric versions of Rad54 and Rdh54 in which the N-terminal domains of these two proteins were swapped. These studies focused on how swapping the N-terminal domains of Rdh54 and Rad54 affected the organization of Rad54 and Rdh54 on Rad51 filaments and D-loop formation. However, they did not address D-loop turnover. The same chimeric versions of Rdh54 and Rad54 are used here. They are composed of aa 1-260 of Rad54 and aa 260-958 of Rdh54 for ^Rad54N^Rdh54, and aa 1-280 from Rdh54 and aa 281-898 from Rad54 for ^Rdh54N^Rad54. Some important biochemical features of these chimera include that both ^Rad54N^Rdh54 and ^Rdh54N^Rad54 are ATPase active and reflect the activity of their parent ATPase. For example, ^Rdh54N^Rad54 has comparable activity to Rad54 [41]. Here we have used the chimeric mutants ^Rad54N^Rdh54 and ^Rdh54N^Rad54 to monitor the effect of translocase positioning on D-loop turnover (Supplemental Figure. 2A). As has been previously reported, the D-loop formation phase of ^Rad54N^Rdh54 was comparable to that of WT-Rdh54, with peak D-loop formation of 5.2±1.2% and 4.8±0.2%, respectively (Supplemental Figure 2BC). Also consistent with previous work, the ^Rdh54N^Rad54 hybrid protein was severely defective in D-loop formation, with peak D-loop formation of 3.1±0.2% as compared to 8.0±1.8% for the WT Rad54 (Supplemental Figure 2BC). It should also be noted that a severe kinetic delay precedes peak D-loop formation, and ^Rdh54N^Rad54 is compromised in this way as well. This is due to defects in homology search [41].

The D-loop turnover phase (10-45 min) of the reaction for both ^Rad54N^Rdh54 and ^Rdh54N^Rad54 indicate defective D-loop turnover, with 91% and 90% of D-loops remaining after 45 minutes, respectively. For WT-Rdh54, 114% of D-loop signal remains (Supplemental Figure 2BC). We analyzed the slope of the turnover phase of the reaction (10-45 min) to measure the rate of D-loop decay (Supplemental Figure 2D). From this measurement, the WT Rad54 decayed with a slope of -1.3×10^−2^. In contrast, D-loops with ^Rad54N^Rdh54 and ^Rdh54N^Rad54 had turnover decay slopes of 1.7×10^−4^ and -0.7×10^−4^, respectively, roughly two orders of magnitude smaller than for WT Rad54 (Supplemental Figure 2D). Similarly, Rdh54 had a turnover decay slope of only 1.7×10^−4^ (Supplemental Figure 2D). These data indicate that both the identity and position of the motor are critical for D-loop turnover, and the Rad54 motor is required for this process.

After determining that the position swap mutants of Rad54 and Rdh54 were unable to promote D-loop turnover, we next asked whether these mutants could inhibit Rad54 from creating 3’ end accessibility at newly formed D-loops. Our expectation from this experiment was that if the position of translocase binding was important for preventing Rad54 activity, then the ^Rad54N^Rdh54 would fail to inhibit 3’ end accessibility of the D-loop while the ^Rdh54N^Rad54 mutant would prevent the accessibility of the 3’ end. Consistent with this prediction, we observed that ^Rdh54N^Rad54 severely inhibited extended D-loop formation with 0%, 0%, and 1.4±1.2% extended D-loops observed at 20, 30, and 45 min, respectively (Fig 5AB). In contrast, the ^Rad54N^Rdh54 was not able to prevent accessibility of the 3’ end, with 1.4±1.2%, 7.4±2.5%, and 27.0±5.0% of extended D-loops observed at 20, 30, and 45 min, respectively (Fig 5AB). We believe the slight reduction in the extended D-loops formed by ^Rad54N^Rdh54 relative to WT Rad54 was due to competition for the same binding site on Rad51. From this experiment we conclude that the position of Rdh54 binding to Rad51 regulates Rad54 activity at D-loops.

**Figure 5:**
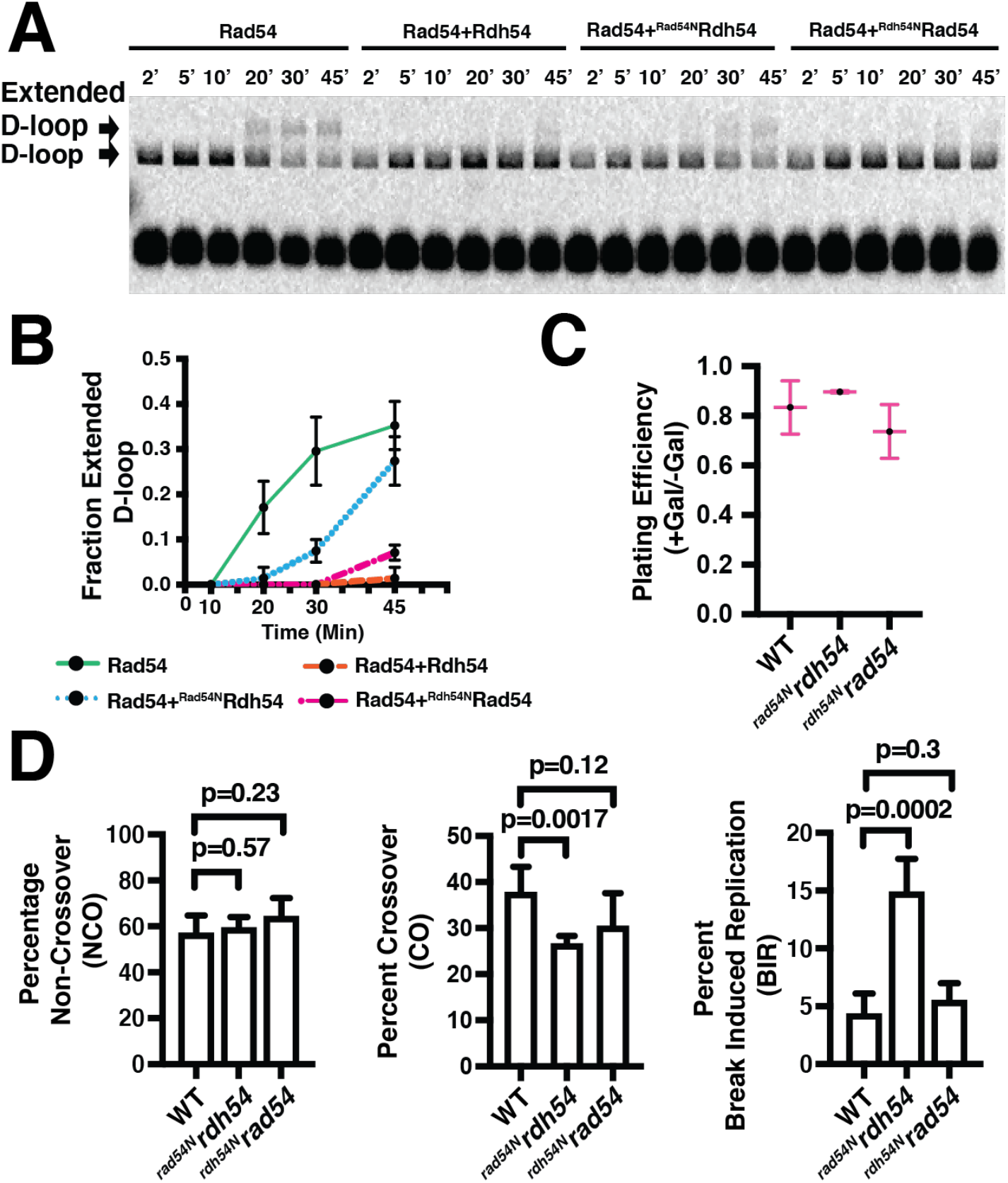
Binding site switching of Rdh54 and Rad54 alters 3’ end accessibility and post gene conversion outcomes. **(A)**. Representative gel image of experiments designed to test how efficiently Rdh54, ^Rad54N^Rdh54, and ^Rdh54N^Rad54 interfere with Rad54 mediated 3’ end accessibility to Klenow fragment. **(B)**. Quantification of extended D-loops for samples with Rad54, Rdh54, ^Rad54N^Rdh54, and ^Rdh54N^Rad54. The data points and error bars represent the mean and standard deviation of at least three independent experiments. **(C)**. Graph representing the plating efficiency (+Gal/-Gal) for WT-*KanMX*, ^*rad54n*^*rdh54*, and ^*rdh54n*^*Rad54* yeast strains. The data represents the mean and standard deviation for four independent experiments. **(D)**. Bar graphs representing the NCO (left), CO (middle), and BIR (right) outcomes for *WT-KanMX*, ^*rad54n*^*rdh54*, and ^*rdh54n*^*rad54*. The data represent the mean and standard deviation of at least four independent experiments. The WT is reproduced from previous figures for comparison purposes.

To genetically test our hypothesis, we integrated the ^*rdh54N*^*rad54* and ^*rad54N*^*rdh54* alleles into the *RDH54* gene, creating diploid strains harboring these mutations and monitored the effect of these N-terminal domain swaps on DNA repair outcomes. From our biochemical observations we expected that the ^*rdh54N*^*rad54* mutant would prevent the onset of BIR but be unable to allow crossovers to occur; a phenotype that resembles *rdh54K318R*. We also expected that the ^*rad54N*^*rdh54* mutant would be unable to prevent BIR outcomes but would be able to promote crossover formation. As with other *RDH54* alleles, there was no significant impact on simple gene conversion (Supplemental Figure 3), and cells survived cleavage by I-SceI (Fig 5C). For ^*rad54N*^*rdh54* allele we observed 26.4±2.2%, 61.0±4.2%, 13.3±2.3% of CO, NCO, and BIR outcomes, respectively (Fig 5D). In contrast, for the ^*rdh54N*^*rad54* allele we observed 30.3±7.2%, 64.1±8.0%, 5.5±1.5% of CO, NCO, and BIR, respectively (Fig 5D). This data indicates that the ^*rad54N*^*rdh54* allele had a 3-fold increase in the amount of BIR compared to the *WT-KanMX* (p-value= 0.0008). From this data we conclude that the position of Rdh54 within Rad51 is critical for complementation of the BIR phenotype.

Surprisingly, the ^*rdh54N*^*rad54* strain showed crossover formation at levels comparable to *RDH54*, suggesting that the Rad54 motor may be able to complement this phenotype through translocase activity. Importantly, this mutation is unable to complement growth sensitivity defects in *rdh54Δ* strain [41]. This suggests that while the prevention of BIR and the ability to allow crossover formation are important for aspects of Rdh54 biology, they do not represent the essential function. As with the other strains analyzed here, there was not a significant loss of chromosomes for ^*rad54N*^*rdh54* or ^*rdh54*^*rad54* (Supplemental Table S1). From this analysis we conclude that the binding position of Rdh54 in D-loops is important for the prevention of BIR, and that the ATPase of Rad54 can partially complement the ability of Rdh54 to open access to the mitotic crossover pathway.

## Conclusion

Here we have gained new insight into the role of Rdh54 during the process of homologous recombination. Our data highlights the ability of Rdh54 to act as a position specific barrier to prevent Rad54 from clearing Rad51 from D-loops. In yeast cells, removal of Rdh54 from its Rad51 binding site results in an increase in gene conversion with BIR. Additionally, an inactive Rdh54 ATPase can still reduce BIR outcomes, but its inability to hydrolyze ATP leads to an increase in NCO outcomes at the expense of CO outcomes. Together, our data supports a model where Rdh54 localization at D-loop intermediates helps regulate genetic exchange post gene conversion and to limit LOH outcomes.

### Rdh54 blocks Rad54 activity at newly formed D-loop intermediates

Physical conflicts between the myriad of proteins present on DNA limit its accessibility and can impede the successful repair of damage during HR. Specific examples of this include the inhibition of Sgs1 translocation by Rad52 [52], the inhibition of Srs2 translocation by Rad54 and Rdh54 [42], and the inhibition of both Srs2 and Sgs1 by the meiosis specific recombinase Dmc1 [53]. Our data indicates that Rdh54 acts as a barrier to Rad54 at newly formed D-loop intermediates, consistent with previous studies that showed Rdh54 limited D-loop size [54]. In contrast to previous reports, we observed that Rdh54 only inhibited the Rad54 mediated D-loop turnover phase of the reaction and did not inhibit D-loop formation.

Our data suggest that Rdh54 protects Rad51 from Rad54 mediated removal at D-loop intermediates. The removal of Rad51 by Rad54 prepares D-loops for DNA extension and promotes DNA template commitment [5, 15, 55, 56]. Rad54 is thought to remove Rad51 by induced ATP hydrolysis of Rad51 in a mechanism resembling the activity of RecA [5, 15, 55, 56]. In contrast, Rdh54 has previously been reported to stabilize Rad51-ssDNA filaments [42], although the exact mechanism is unclear. Several possibilities include the stabilization of Rad51 against ATP hydrolysis [42], the physical interference of Rad54 translocation, or by limiting Rad54 movement through the interaction between Rdh54 and Rad51. Although somewhat controversial, a link between Rdh54 and locating donor DNA has also been reported using a donor less strain of yeast [57, 58], suggesting stabilization of Rad51 filaments could have a limited role in the process of locating the correct donor DNA molecule. For example, Rdh54 stabilization of Rad51 filaments may prevent Rad54 from destabilizing Rad51 at DNA sites that contain limited homology, improving the fidelity of homology search; however, this idea remains to be tested.

While our *in vitro* assays illustrate limited accessibility of the 3’ end of DNA at newly formed D-loops, the Rdh54 mediated block must be removed *in vivo* to allow gene conversion. In cells it is likely that the Rdh54 barrier is only temporary, and acts to delay clearance of proteins from the 3’ end of DNA. This possibility leads to several ways in which inhibition could be relieved. First, Rdh54 translocation or dissociation away from Rad51 at D-loops may allow Rad54 to remove Rad51. A second possibility is that spatial separation between Rdh54 and Rad54 regulates how much Rad51 is removed from D-loops. In this scenario Rdh54 may not prevent disruption of the 3’ end but may prevent complete disruption of Rad51 recombinase filaments. Alternatively, a combination of these two possibilities may regulate the position and activity of a Rdh54 barrier and the accessibility of the 3’ end of DNA as HR proceeds. Further work, including measuring the frequency of Rad54 and Rdh54 colocalization at D-loops *in vivo*, will be required to determine how Rdh54 inhibition of Rad54 mediated Rad51 disruption is relieved. Despite these open questions, our observation that *RDH54* does not affect gene conversion but rather the extent of genetic exchange between donor and recipient DNA molecules indicates that stabilization of Rad51 filaments is likely to be important for the transfer of information between DNA molecules.

### The implications of Rdh54 positioning during HR

As previously reported, the Rad51 presynaptic complex maintains two sites for active translocase binding. Rdh54 fails to promote D-loop turnover and accessibility of the 3’ end of DNA even when moved to the binding position of Rad54. This data indicates that Rdh54 is unable to remove Rad51 from D-loops or promote template commitment like Rad54 and suggests that Rdh54 may occupy a secondary recombination site that likely remains active during strand invasion through stabilization of Rad51. Interestingly, the ^*rdh54N*^*rad54* allele was genetically sufficient to complement both phenotypes observed for Rdh54, suggesting that an ATPase in the correct position was sufficient to carry out function during HR. However, this allele is unable to complement DNA damage sensitivity phenotypes associated with *rdh54*Δ cells [41], and it is likely that *RDH54* has another specific role during DNA damage response in cells which may be linked to cellular recovery [60].

Our data also suggests that improper positioning of Rdh54 on Rad51 filaments, or deletion of *RDH54*, results in an increase in BIR outcomes [29, 49]. A tentative interpretation of this data is that *RDH54* may reduce the on-set of BIR by slowing the accessibility of the DNA 3’ end preventing premature initiation of DNA repair (Fig. 6A). One possible way this could affect HR outcomes is to allow both ends of the DSB to synchronize their size and organization to allow effective annealing and subsequent repair of both ends through SDSA or DSBR [29]. Thus, the presence of Rdh54 would favor SDSA/DSBR over BIR. One drawback of this model is that expanded D-loops are a preferred substrate for second end capture in the DSBR model [20-22]. Therefore, delaying D-loop expansion may also limit the annealing of the second end of DNA to D-loops and ultimately favor SDSA, which may help explain why crossover outcomes are lost in *rdh54K318R* mutants (see below).

**Figure 6:**
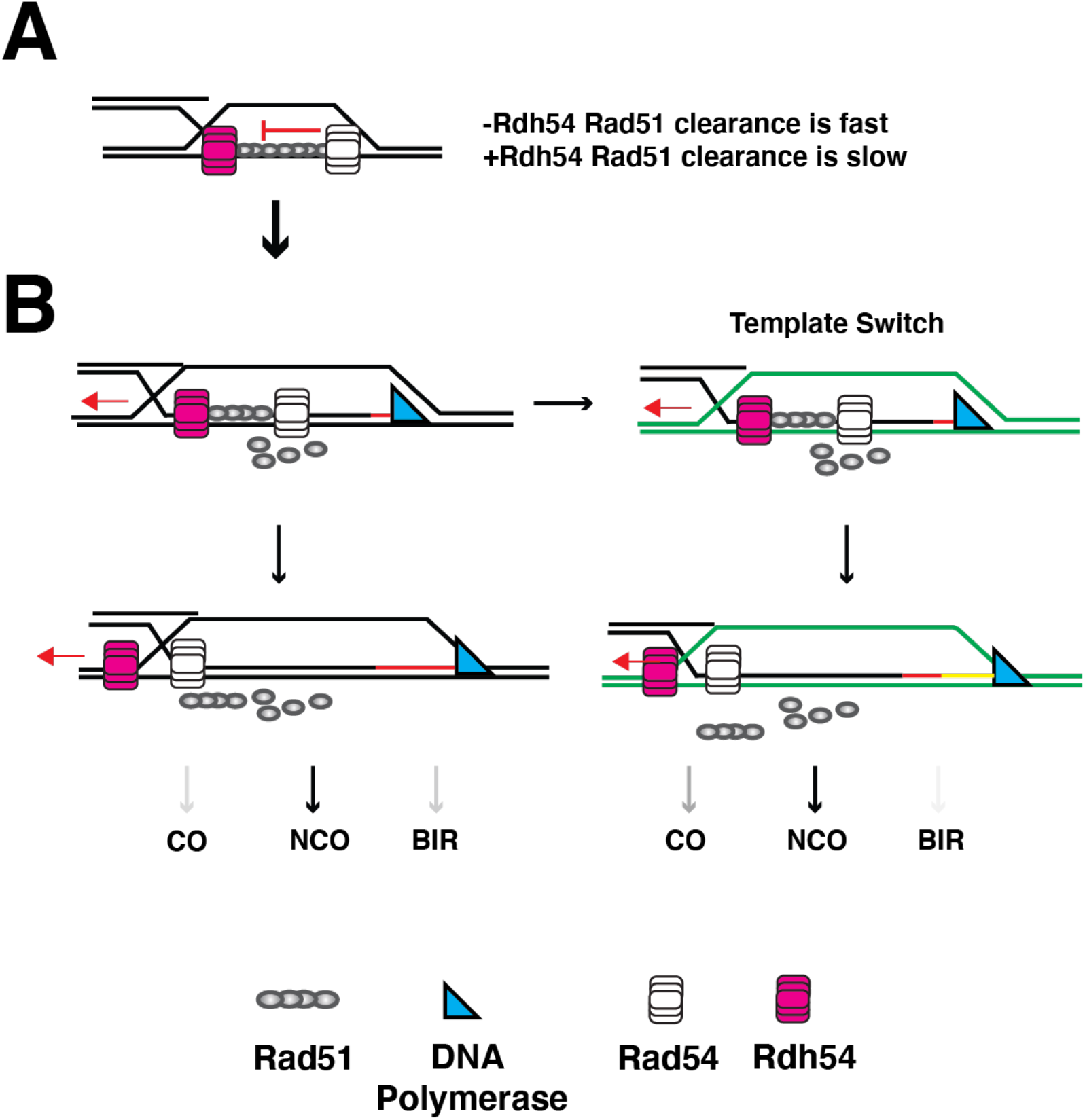
Model for how Rdh54 alters HR pathway choice. **(A)**. A diagram illustrating the first step in our model for Rdh54 regulation of D-loop intermediates. When Rdh54 is absent the removal of Rad51 is fast. When Rdh54 is present the removal is slow, changing the rate of template choice. **(B)**. Rdh54 maintains the ability of D-loops to undergo template switching and eventually leaves the D-loop. This leads to a reduction in BIR outcomes and ultimately a change in the distribution of CO, NCO, and BIR outcomes. The change in distribution is highlighted by differential shading of the arrows.

Previous reports have not identified an active role for *RDH54* in inhibition of *RAD51* mediated BIR. However, *RDH54* has been implicated in ICTS [7, 59], a type of DNA template switching that occurs post initiation of BIR. ICTS can result in the synthesis of DNA from multiple homologous DNA sequences and in diploids can create mosaic patches of sequence from homologous chromosomes [59]. Considering these observations, an alternative interpretation of our data is that deletion of *RDH54* results in loss of ICTS. This would lead to an increase in chromosomal template commitment through Rad54-mediated displacement of Rad51, and an apparent increase in long range template committed BIR. Our reasoning behind this possible model is that in the genetic assay used in this study, BIR must occur over at least 154 kb for the appropriate marker to be copied. Template switching prior to copying the marker will prevent the observation of BIR outcomes. Therefore, the observed increase in BIR could also be the result of a failure to switch chromosomal templates. Interestingly, it has previously been shown that loss of Rdh54 catalytic activity is required for its role in ICTS [7]. In contrast, our BIR phenotype is complemented with the catalytically inactive *rdh54K318R* allele. However, in a situation where the homologous chromsomes are not divergent, stabilization of Rad51 filaments may be sufficient to promote template switching. Therefore, this observation does not effectively differentiate between these two models. Despite their mechanistic differences, in both models *RDH54* regulates the transfer of information between DNA molecules.

### Access to crossover pathways

We observed that the *rdh54K318R* allele resulted in an increase in NCO and a reduction in CO outcomes. This observation is consistent with recent findings that *rdh54K318R* showed a reduction in crossovers, as well as a kinetic delay in general repair in haploid yeast [9]. These findings likely indicate that Rdh54 must move or translocate away from sites of recombination in order for gene conversion with crossovers to occur. This does not mean *RDH54* is required for crossovers, as crossovers form normally in *rdh54Δ* strains. Although the *rdh54K318R* allele is a permanent case of translocase inactivation that does not occur in cells, temporary inactivation *in vivo* may modulate the ATPase of Rdh54. If Rdh54 were held inactive it would promote NCO outcomes, preventing the LOH through crossover outcomes. Recently, a role for Rdh54 has been observed for D-loop expansion [9], a process involving template commitment by Rad54 and DNA synthesis by DNA polymerase delta [1]. Our data suggests that Rdh54 may regulate this process through stabilization of Rad51 within D-loops. ATP hydrolysis may be required to access the crossover pathway by removing Rdh54 and allowing complete removal of Rad51 or granting access of Rad54 to the DNA junctions promoting branch migration [61]. Alternatively, pathways such as SDSA that promote NCO outcomes may be favored by incomplete clearance of Rad51 from the D-loop.

A general hypothesis from our data (Fig 6AB) is that upon initial strand invasion Rad54 clearance of Rad51 is fast in the absence of Rdh54, and slow in the presence of Rdh54. This reduces the rate of template commitment and can allow ICTS by maintaining active Rad51. Over time, Rdh54 can translocate or be removed from strand invasion intermediates allowing template commitment and complete repair of DNA. When Rdh54 is absent, BIR or BIR with template switching is high, and when Rdh54 is inactive NCO are high. Therefore, the presence of Rdh54 regulates the amount of genetic exchange that can occur between chromosomes (Fig 6AB). Despite the precise mechanisms behind the function of Rdh54, a general conclusion in the context of our study is that Rdh54 can function to reduce LOH.

### Conservation of a two-paralog system in higher eukaryotes

We used a combination of genetic and biochemical experiments to show that the *S. cerevisiae* DNA motor protein Rdh54 acts as a regulator of genetic exchange between donor and recipient DNA molecules during HR. However, a two Rad54 paralog system is also conserved in most eukaryotes. For example, in higher eukaryotes Rdh54 is believed to be conserved as RAD54B [62], and Rad54 is conserved as RAD54L [63]. Currently, it remains unclear if RAD54B is a direct homolog of Rdh54 or Rad54. This leads to the possibility that RAD54B may only be functionally homologous to Rdh54. Along these lines, RAD54B shares several functional similarities to Rdh54, including roles in cell cycle recovery from DNA damage and general biochemical activities [60, 64, 65]. Future work will be directed at understanding if the regulation of genetic exchange by Rdh54 is conserved in higher eukaryotes, how movement of Rdh54 relieves the barrier to mitotic crossover formation, and the precise mechanisms of Rdh54 activity during ICTS.

## Materials and Methods

### Protein Purification

Purification of Rad51, RPA, GST-Rad54, and GST-Rdh54 were carried out as previously described [41, 56, 66, 67].

### Yeast Strain generation

Gene deletions were generated by PCR amplification of *KanMX* selectable marker with 50 nt of homology upstream and downstream of the ORF of the *RDH54* gene and transformed into WT-15D with selection on YPD+G418 (500 µg/ml). Genetic knockouts were confirmed by amplification of yeast genomic DNA with primers that annealed 250 bp upstream or downstream of the *RDH54* ORF. Subsequent deletions were generated by amplification of the region of the genome and transforming WT-11C. A similar strategy was used for *RAD54* deletions. For genetic knock-ins, a *KanMX* marker was integrated 125 bp downstream of the *RDH54* stop codon in a plasmid containing the *RDH54* gene. *WT-KanMX* and *rdh54K318R-KanMX* were PCR amplified and used to replace the endogenous *RDH54* locus in both WT-11C and WT-15D. Integration was confirmed by PCR. Full strain genotypes are in SI Appendix Table 2.

### Red/White homologous recombination assay

The WT strains used in this assay as well as the procedure for diploid formation are described here [47, 48]. The genotypes for modifications to these strains can be found in SI Appendix Table 2. The assay was performed by growing the appropriate strain overnight in YP +2% raffinose. The next day cells were diluted to an OD_600_ of 0.2 and allowed to reach an OD_600_ of 0.4-0.5 followed by the expression of I-SceI by the addition of 2% galactose. Cells were allowed to grow for an additional 1.5 hours after which they were plated on YP+2% dextrose and allowed to grow for 48 hours. After 48 hours they were placed in the 4°C overnight to allow further development of red color. The number of white, red, and sectored colonies was then counted followed by replica plating onto YPD+ hygromycin B (200 µg/ml) and YPD+ nourseothricin (67 µg/ml, clonNat) for analysis of recombination outcomes. Technical duplicate plates were also plated onto YNB (-ade) +2% galactose to measure reinduction of I-Sce1, indicating the fraction of cleaved sister chromatids. Finally, strains were also plated on YNB (-URA/-MET) + 2% dextrose to insure proper chromosome segregation. The data was analyzed by counting colonies that were red, white, or sectored, and then counting sectored colony survival on different antibiotic sensitivities. The data for each category was then divided by the total population of sectored colonies. The standard deviation between biological replicates analyzed for at least three independent experiments.

### D-loop Assay

D-loop formation experiments were performed in HR buffer (30 mM Tris-OAc [pH 7.5], 50 mM NaCl, 10 mM MgOAc_2_, 1 mM DTT, 0.2 mg/ml BSA) using an Atto647N labeled DNA duplex (15 nM) consisting of 21 nt overhang that was homologous to a region on the pUC19 plasmid. Additionally, some experiments were performed with an Atto647N labeled 90 mer ssDNA that was homologous to the pUC19 plasmid. Rad51 (300 nM) was incubated with recipient DNA at 30 °C for 15 minutes. The resulting Rad51 PSCs were then mixed with indicated concentrations of Rad54, Rdh54 or Both, RPA (500 nM) and pUC19 plasmid (0.3 nM). Reactions were quenched after indicated periods of time.

### 3’ end extension assay

Native 3’ end extension assays experiments where set-up as describe in the D-loop assays with the exception that dNTP’s (1 mM) and Klenow (exo-)(NEB) were added after 10 minutes of D-loop formation. For denaturing 3’ end extension assays experiments were set-up using the same conditions and concentrations as D-loop assays with the exception that after 10 min dNTPs and Klenow (exo-) were added and allowed to incubate at 30°C. The reactions were quenched as described above and incubated with 1 unit of Proteinase K for 37°C for 20 minutes. The reactions were then resolved by electrophoresis on a 0.9% agarose gel and imaged for fluorescence.

### ATP hydrolysis assay

ATP hydrolysis assays were performed in HR buffer (30 mM Tris–OAc [pH 7.5], 50 mM NaCl, 20 mM MgOAc, 1 mM DTT, 0.2 mg/ml BSA) in the presence of 1 mM ATP and trace amounts of γ^32^P–ATP. All reactions were performed at 30°C and contained pUC19 plasmid dsDNA (50 ng/µl). Aliquots were removed at specified time points and quenched with 25 mM EDTA and 1% SDS. The quenched reactions were spotted on TLC plates (Millipore, Cat. No. HX71732079) and resolved in 0.5 M LiCl plus 1 M formic acid. Dried TLC plates were exposed to phosphorimaging screen and scanned with a GE Healthcare Life Sciences Typhoon FLA 9500 biomolecular imaging system.

### Total protein measurement

Total protein measurements were made as described in Silva et al. [51]. Briefly, *RAD54-YFP* (strain ML1067) or *RDH54-YFP* (strain ML906) cells were estimated by measuring total nuclear YFP fluorescence intensities using Volocity software.

## Acknowledgements

Lorraine Symington for providing the original yeast strains. We thank Eric Alani for plasmids and helpful comments and discussion, and Eric Greene, Marcus Smolka, and Rodney Rothstein for helpful comments and discussion. We also thank members of the Crickard laboratory for critical reading of the manuscript. This work is funded in part by NIH grants R35GM142457 to JBC. ML was supported by Independent Research Fund Denmark (research grant 1026-00041B) and the Villum Foundation (research grant 11407).

## Author Contributions

M.K. conducted and analyzed all genetic experiments. J.D. contributed reagents. ML performed molecules per cell experiments. J.B.C performed protein purification, biochemical experiments, and prepared the manuscript with input from M.K., J.D, BHF, and ML.

## Supplemental Information

**Supplemental Table S1.** All red/white outcomes CO/NCO/BIR.

All red/white outcomes CO/NCO/BIR
| Strain | Red | White | Sectored | Total | CO | NCO | BIR | Chromosome loss |
| --- | --- | --- | --- | --- | --- | --- | --- | --- |
| WT | 457 | 194 | 960 | 1611 | 398 | 504 | 58 | 1/145 |
| *WT-KanMX/WT-KanMX* | 307 | 50 | 548 | 905 | 200 | 323 | 25 | 0/325 |
| *RDH54/rdh54Δ* | 233 | 52 | 339 | 624 | 140 | 163 | 36 | 0/223 |
| *rdh54Δ/rdh54Δ* | 238 | 97 | 301 | 636 | 104 | 145 | 52 | 0/67 |
| *rad54Δ/rad54Δ* | 31 | 0 | 0 | 31 | N/A | N/A | N/A | N/A |
| *rdh54K318R/rdh54K318R* | 285 | 21 | 381 | 687 | 56 | 311 | 14 | 0/381 |
| *rdh54<sup>N</sup>Rad54/rdh54<sup>N</sup>Rad54* | 568 | 56 | 936 | 1560 | 286 | 626 | 51 | 0/324 |
| *rad54<sup>N</sup>Rdh54/rad54<sup>N</sup>Rdh54* | 453 | 60 | 632 | 1145 | 166 | 381 | 85 | 0/79 |

**Supplemental Table S2.** Yeast Strains used in this study.

Yeast Strains used in this study
| Strains | Genotype* | Source or reference |
| --- | --- | --- |
| LSY-2202-15D (WT) | *MATa ade2-n his3::NatMX4 met22::klURA3* | (Mazon et al. 2010) |
| LSY-2205-11C (WT) | *MATalpha ade2-I lys2::GAL-ISCEI his3::HphMX4* | (Mazon et al. 2010) |
| JBC001 (15D) | *rdh54Δ::KanMX* | This study |
| JBC002 (11C) | *rdh54Δ::KanMX* | This study |
| JBC007 (11C) | *rad54Δ::KanMX* | This study |
| JBC010 (15D) | *rad54Δ::KanMX* | This study |
| JBC0011 (15D) | *RDH54::RDH54-KanMX* | This study |
| JBC0013 (11C) | *RDH54::RDH54-KanMX* | This study |
| JBC0014 (15D) | *RDH54::rdh54K318R-KanMX* | This study |
| JBC0016 (11C) | *RDH54::rdh54K318R-KanMX* | This study |
| JBC0039 (15D) | *RDH54::rdh54<sup>N</sup>RAD54-KanMX* | This study |
| JBC0041 (11C) | *RDH54::rdh54<sup>N</sup>RAD54-KanMX* | This study |
| JBC0043 (15D) | *RDH54::rad54<sup>N</sup>RDH54-KanMX* | This study |
| JBC0045 (11C) | *RDH54::rad54<sup>N</sup>RDH54-KanMX* | This study |

## Supplemental Figures and Legends

**Supplemental Figure 1:**
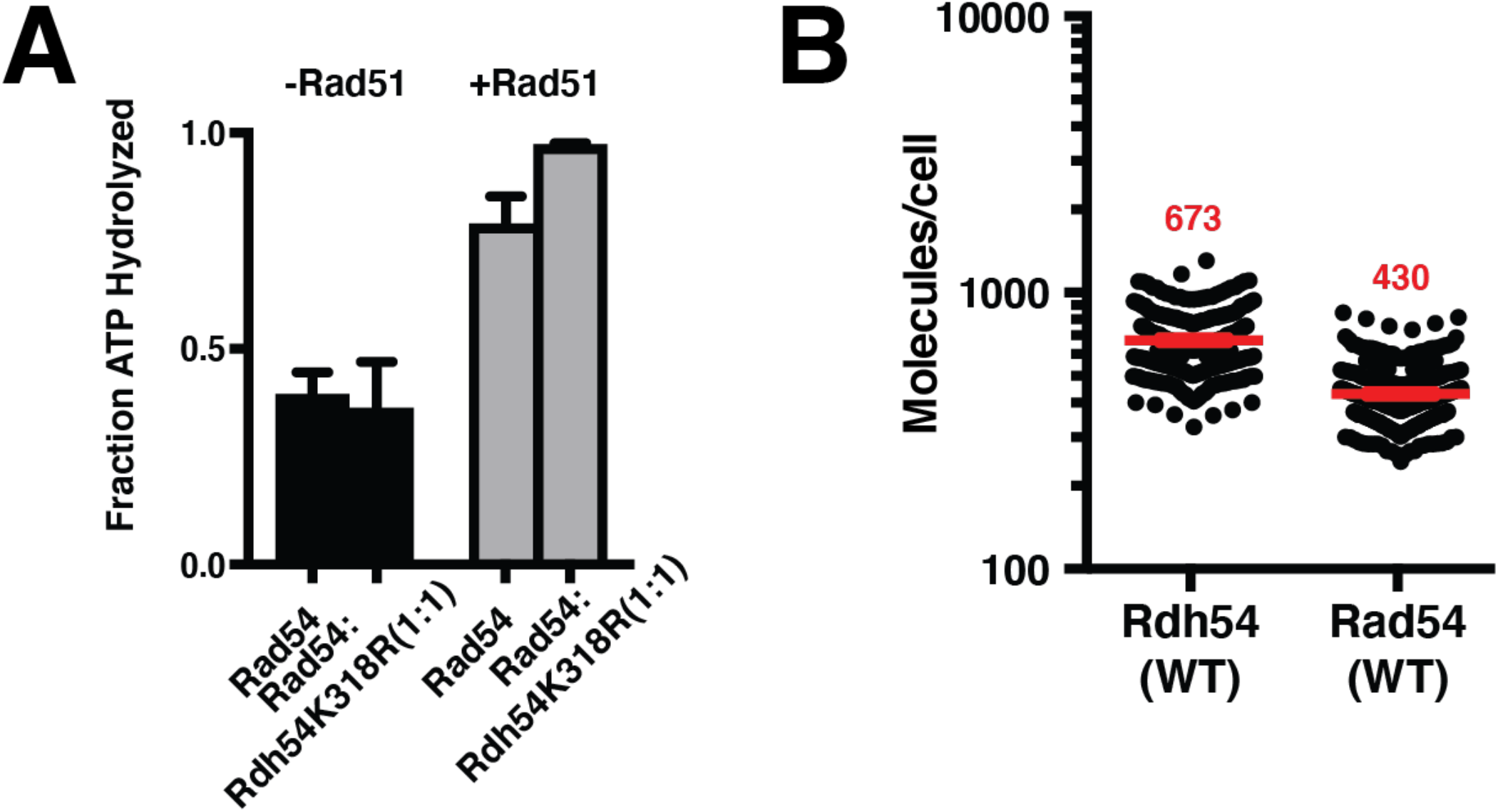
Rdh54 does not alter Rad54 ATPase functions at physiological ratios. **(A)**. ATPase assay illustrating the fraction of Rad54 ATP hydrolysis in the presence and absence of Rad51 with and without Rdh54K318R. The bars represent the mean and standard deviation of three independent experiments. **(B)**. Quantification of the number of fluorescently labeled Rad54 (N=100) and Rdh54 (N=100) molecules in yeast cells. The line and error bars represent the mean and 95% confidence interval of the data. The red number above the data is the mean molecules per cell.

**Supplemental Figure 2:**
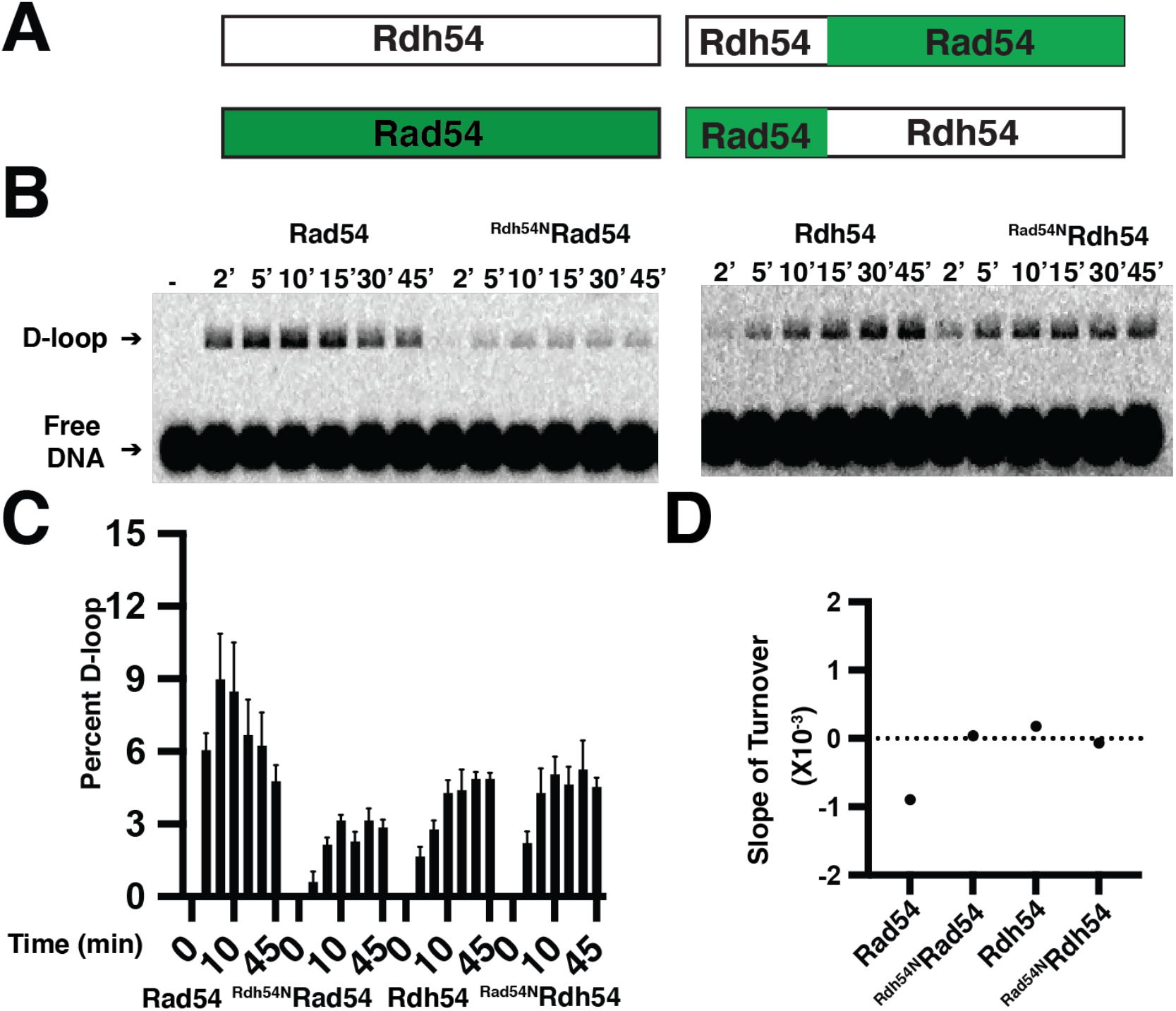
N-terminal chimeras of Rad54 and Rdh54 show differences in D-loop turnover. **(A)**. Schematic illustrating domain swap constructs for ^Rdh54N^Rad54, and ^Rad54N^Rdh54. These mutants alter the position of translocase binding on Rad51 filaments. **(B)**. Representative agarose gel illustrating the formation and disruption of D-loops in the presence of Rad54 (left), ^Rdh54N^Rad54 (middle left), Rdh54 (middle right), ^Rad54N^Rdh54 (right). **(C)**. Quantification of D-loop formation and disruption in the presence of Rad54 (left), ^Rdh54N^Rad54 (middle left), Rdh54 (middle right), Rad54NRdh54 (right). The columns and error bars represent the mean and standard deviation of three independent experiments.

**Supplemental Figure 3:**
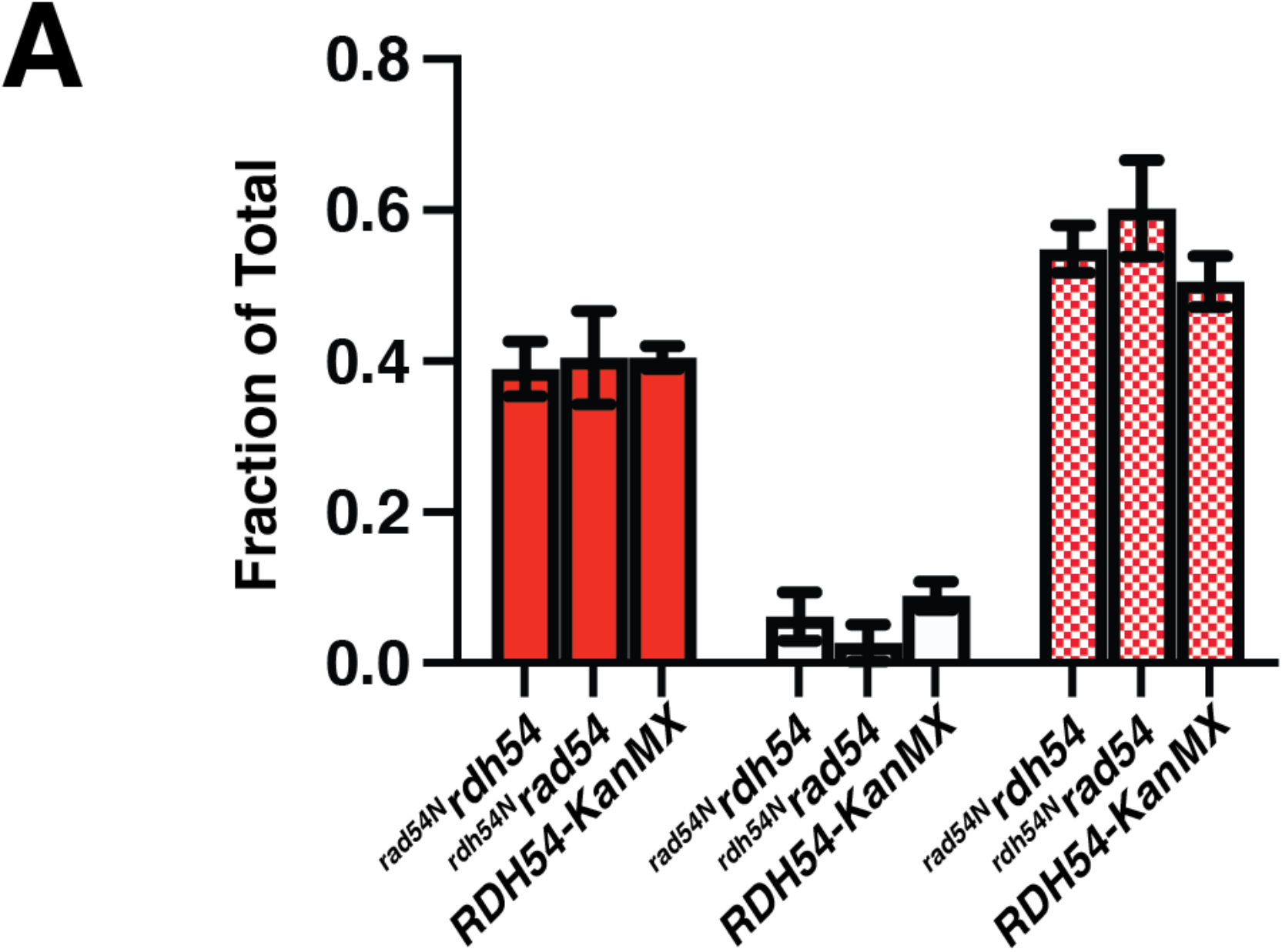
Gene conversion outcomes are unaffected in N-terminal chimeric Rad54 and Rdh54. **(A)** Graph representing the red/white/sectored colony outcomes for all strains used in this paper. Strains are labeled in the figure. The bars and error bars represent the mean and standard deviation of at least three independent experiments.

